# Deferred mortality: cyclic thermal stress during pupation triggers irreversible carry-over effects in a key pollinator

**DOI:** 10.64898/2025.12.07.692821

**Authors:** Maximilian M. Mandlinger, Christoph Kurze

## Abstract

Extreme heatwaves threaten global biodiversity, yet the sublethal carry-over effects of developmental thermal stress on adult fitness remain poorly understood. We addressed this knowledge gap for bumblebees, key pollinators that face alarming population declines, using a controlled *in vitro* approach. We exposed *Bombus terrestris* pupae to cyclic thermal stress to evaluate emergence success, adult longevity, and morphological traits. Cyclic thermal stress caused significant acute pupal mortality (up to 30% reduced emergence) and, crucially, deferred mortality, significantly reducing adult longevity (hazard ratio, HR = 2.00) in both workers and males. Rearing *in vitro* at 34°C showed no significant difference compared to individuals that developed naturally in their natal colony (HR = 0.95). Antennal and wing deformations served as powerful hazard indicators (HR = 2.38) for premature adult mortality. Our findings reveal that cyclic thermal stress during pupal development imposes irreversible developmental damage, undermining adult physiological resilience.

## 1. Introduction

Extreme weather events, such as heatwaves, are increasing in frequency and intensity due to climate change, leading to distinct and widespread implications for biodiversity, including insects [1, 2]. While thermal stress is known to have lethal and sublethal effects across a wide range of insects, most studies have focused on acute effects in adults, such as mortality at critical thermal maxima, metabolic disruption, and impaired flight performance and cognition [3-5]. Little is known about how thermal stress during an earlier life stage impose sublethal, yet long-lasting, carry-over effects on adult fitness [6-9]. Gaining a better understanding of such sublethal carry-over effects, impacting colony fitness, is essential for accurately forecasting population declines in keystone insect taxa such as bumblebees (*Bombus* spp., Hymenoptera: Apidae) [10].

Bumblebees, as a cold-adapted taxon, have experienced population declines, which are partially attributed to climate change [11-16]. This is a concerning trend, given that they provide crucial pollination services in ecosystems and agriculture [17, 18]. While ground-nesting species can mitigate extreme ambient temperatures to some extent, other species, such as *B. hypnorum* and *B. pascuorum* that typically nest above the ground [19], or commercially used species in greenhouses such as *B. terrestris* [20], face a higher risk of thermal stress. Colony-level studies have demonstrated that constant prolonged elevated temperatures negatively impact colony energetics, growth and reproductive fitness [21-24], and such thermal stress has been linked to altered offspring morphology [25-27]. As behavioural responses vary widely between colonies, it is difficult to disentangle indirect effects arising from worker thermoregulation [28, 29], from direct physiological effects [30]. For example, if workers need to spend more time fanning to cool the nest or incubating the brood to maintain optimal brood temperatures (generally above ambient nest temperatures) rather than foraging and nursing [28, 29], this can affect larval development and ultimately colony fitness. To gain a precise mechanistic understanding of how thermal stress affects specific larval stages, *in vitro* rearing offers the required control at the cost of losing natural realism [30, 31].

Our previous *in vitro* studies have already revealed stage-specific acute responses to constant thermal stress in *B. terrestris* larvae and pupae [30, 31]. While smaller fourthinstar (L4) larvae may have a selective advantage under thermal stress due to their ability to more easily reach the critical body mass for pupation [31], thermal stress at the pupal stage appears to select for individuals with increased relative lipid reserves [30]. A constant *in vitro* rearing temperature of 38°C during L4 larval and pupal development was identified as a critical thermal threshold, resulting in an increased direct mortality and higher a incidence of wing deformations [30, 31]. While these studies illustrate impaired larval and pupal development at constant thermal stress, the magnitude of the downstream impact on adult survival and the effect of fluctuating thermal stress scenarios remains unknown.

To address this fundamental knowledge gap, we used highly controlled *in vitro* rearing of *B. terrestris* pupae to isolate the effect of cyclic thermal stress scenarios on individual life history. Our main aim was to assess the carry-over effects under fluctuating thermal extremes, an approach more ecologically relevant than constant temperature regimes [21, 30, 32, 33]. We focused on the pupal stage as metamorphosis in holometabolous insects, such as bees, represents the most extreme transition between phenotypes [34] and is likely the most vulnerable to environmental stress [30, 35]. We quantified both the direct effect on emergence success and the subsequent long-term adult survival, morphology, and energy reserves. We hypothesised that increasing cyclic thermal stress during pupal development (metamorphosis) would affect adult morphology (specifically wing morphology [30, 35]) and trigger deferred mortality, significantly reducing adult longevity. Furthermore, we expected that the impact of thermal stress would differ between workers and males.

## 2. Materials and methods

### 2.1 Colony husbandry

We maintained ten queen-right source colonies of *B. terrestris* (Natupol Research Hives, Koppert B.V., Netherlands) under standardized laboratory conditions [36]. Colonies were housed in standard nest boxes (23 × 25 × 15 cm), which were connected to foraging arenas (59 × 39 × 26 cm). We maintained all colonies at 25 ± 1°C and 30-50% RH, under a 14:10 h light:dark regime. Bumblebees had *ad libitum* access to a 66% w/w sucrose solution (Natupol Bee Happy, Koppert B.V., Netherlands). We supplied each colony with 6-11 g of pollen candy (67% w/w organic pollen, 25% w/w sucrose, 8% w/w water) daily.

### 2.2 Larvae collection and *in vitro* rearing

We collected 4^th^ instar (L4) larvae from fully developed source colonies (for sample size see Table 1; Fig. S1*a*). Collecting L4 larvae was crucial to ensure that we could start all thermal regimes consistently at the beginning of the pupal phase. Larvae were reared *in vitro* following previous protocols [for details see 30, 31, 37].

**Table 1.** Overview of the experimental design. This table details the rearing conditions (regime and temperature) of pupae for all treatment groups, including individually developed naturally in their natal colony. Sample sizes (n) reflect the total number of pupae per treatment at start of *in vitro* rearing experiment (indicated by ^§^) and emerged adults at the start of the survival assay (indicated by ^#^). N indicates the number of colonies used in each treatment group.

| Treatment | Regime<sup>§</sup> | Temperature<sup>§</sup> | Sample size (n) | N |
| --- | --- | --- | --- | --- |
| colony-reared | developed naturally | $25 \pm 1^\circ\text{C}^*$ | <sup>#</sup>115 | 10 |
| 34°C | constant | 34°C | <sup>§</sup>132/ <sup>#</sup>102 | 6 |
| 37°C cyclic | 12:12 h cycle | 37°C:34°C | <sup>§</sup>125/ <sup>#</sup>82 | 5 |
| 38°C cyclic | 12:12 h cycle | 38°C:34°C | <sup>§</sup>146/ <sup>#</sup>78 | 5 |

\* colonies kept in the insectary
<sup>§</sup> pupae / <sup>#</sup> adults

Briefly, L4 larvae were carefully transferred into 3D-printed polylactide (PLA) artificial brood cells (Fig. S1*b*), weighed (initial larval weight; d = 0.1 mg, Sartorius M-Pact AX224, Sartorius GmbH, Germany), then placed inside 24- or 48-well clear flat bottom plates (Fig. S1*c*; Falcon/Corning, USA) and put within a humidity-controlled plastic container (18.5 × 18.5 × 11.5 cm) kept at a constant temperature of 34°C (KB115, BINDER GmbH, Germany). While 34°C is at the upper range of the typical nest temperature in *B. terrestris* [19], it aligns with brood temperatures, which are typically 2°C warmer than ambient nest temperatures [28] and likely represent the optimal temperature for larval development [29]. Moreover, 34°C has been used in previous *in vitro* rearing protocols for *B. terrestris* [30, 31, 37], making it comparable to previous *in vitro* studies. To maintain a relative humidity (RH) of 65 ± 10%, a 120 mL cup of saturated sodium chloride solution was also placed within the container.

Larvae were fed a pollen medium (Fig. S1*b*), consisting of 50% w/v sucrose solution, 40% organic pollen, 10% yeast extract (Bacto™, BD, USA), and 1% casein sodium salt from bovine milk (Sigma-Aldrich, Germany) [37], twice daily to satiation [31]. Feeding was stopped with the beginning of the pupal phase. Our setup allowed us to measure pupal weight non-invasively and facilitated the easy removal of any dead individuals.

### 2.3 Experimental procedure

At the beginning of pupation (metamorphosis), each individual was weighed (initial pupal weight; Sartorius M-Pact AX224) and randomly assigned to one of three experimental treatment groups (for sample sizes see Table 1). Pupae were transferred to new, humidity-controlled containers within one of three programmed incubators. The control group was maintained at a constant temperature of 34°C (34°C, constant *in vitro* control). The two cyclic thermal stress treatment groups experienced a 12:12 h temperature cycle between a baseline of 34°C (night phase) and a thermal maximum (day phase) of either 37°C (37°C cyclic) or 38°C (38°C cyclic) until adult emergence (i.e. 9-10 days for males, 8-9 days for workers; Fig. 1*c*). Given that bumblebee species regulate their core nest temperatures between 30-35°C for optimal brood development [19], our two cyclic temperature treatments simulate a thermal stress scenario when colony thermoregulation fails during the day. Pupal survival and adult emergence were monitored blindly twice daily. In addition to our *in vitro* treatments, newly emerged adults (< 1 day old) were collected directly from their natal colonies to serve as an additional control group (colonyreared) for the subsequent longevity assay.

**Figure 1.**
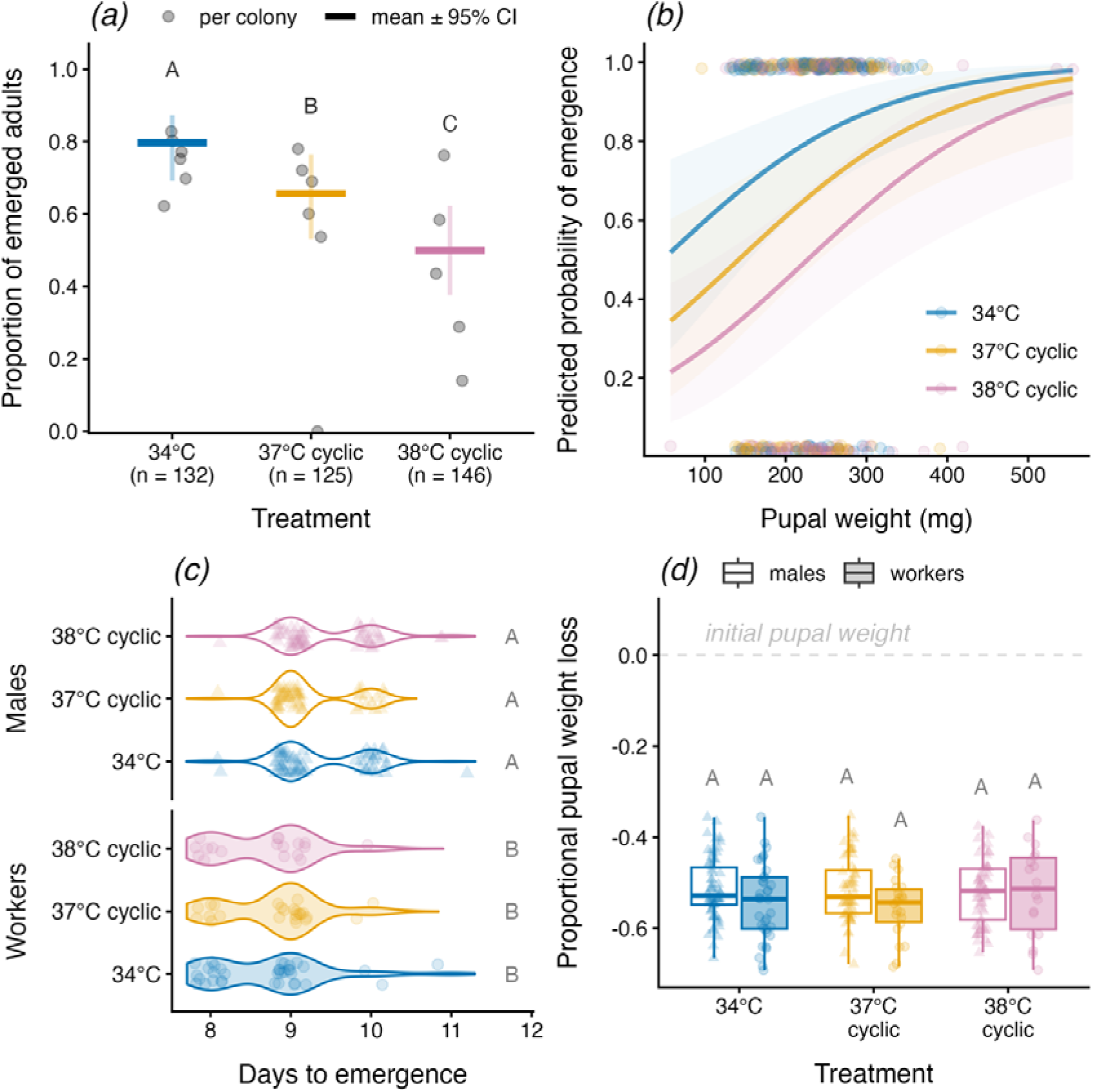
Direct effects of cyclic thermal stress during pupation on emergence success and development duration in *B. terrestris*. ***(a)*** Proportions of pupae emerging as adult bees observed per colony (grey points; n ≥ 5 per colony) and GLMM means ± 95% CI (bars). Sample sizes (n) across all colonies are provided below for each treatment group. ***(b)*** Predicted probability of emergence (± 95% CI, shaded areas) in relation to pupal weight and treatment. ***(c)*** Pupation duration until emergence for males (open violins) and workers (filled violins). **(*d*)** Proportional pupal weight loss from the beginning of pupation until emergence as adult males (open boxes) and workers (filled boxes). Colours **(*a-d*)** indicate different *in vitro* rearing conditions for pupae reared at constant 34°C (blue), and under cyclic thermal stress regimes of 37°C (orange) and 38°C (reddish-purple). Individual data points in **(*b-d*)** are shown as triangles (males, m) and circles (workers, w). Sex-specific samples sizes for 34°C (n_m_ = 67, n_w_= 35), 37°C cyclic (n_m_ = 58, n_w_ = 24), and 38°C cyclic (n_m_ = 60, n_w_ = 18). Significant differences (p < 0.05), based on post-hoc comparisons from GLMMs, are indicated by different letters above the bars and next to the violin plots (groups sharing a letter are not significantly different).

### 2.4 Adult survival assay

Newly emerged adult bees from each thermal treatment (i.e. those that experienced as pupae *in vitro*) and the additional colony-reared group (for sample sizes, see Table 1) were weighed (emergence weight; Sartorius M-Pact AX224) and then individually housed in modified 50 mL tubes (Fig. S1*d*; Greiner Bio-One GmbH, Germany). These were stored in a wooden rack within the insectary room housing their natal colonies at 25 ± 1°C and 30-50% RH (Fig. S1*e*). Each tube contained a 3D-printed PLA mesh insert for sanitary reasons. The lids were modified to include three 3 mm ventilation holes and one 1 cm feeding port, through which a 1.5 mL perforated tube provided *ad libitum* access to a 60% w/v sucrose solution. Pollen candy was offered separately in the caps of 2 mL reaction tubes and was replaced weekly or as necessary. Housing tubes and meshes were replaced as needed to ensure sanitary conditions. Adult survival was monitored around noon daily. Dead individuals were recorded and immediately frozen at -21°C until further analysis.

### 2.5 Morphological and physiological analysis

We conducted the morphological and physiological analyses on adult workers and males, which were exposed to different thermal scenarios as pupae (Table 1), following our previous protocols [30, 31, 36]. In addition to their emergence weight, the intertegular distance (ITD) and head width, used as proxies for adult body size, were measured on dead bees from the survival assay using a digital microscope (VHX-500F, Keyence GmbH, Germany). The sex of each adult bee was determined by counting flagellomeres (workers = 10; males = 11). Furthermore, any deformations of their antennae were recorded (Fig. S2*a*). To assess wing integrity, forewings were carefully detached, embedded side-by-side on a microscope slide (Roti^®^Mount, Carl Roth GmbH + Co. KG, Germany) and photographed at 20× magnification. Wings were subsequently classified as deformed (figure S2*b*) when visually apparent malformations were present.

The remaining bee bodies were processed to determine dry mass and relative lipid content following our previous protocol. Specimens were first cut open from the stinger to the fourth sternite and then dried at 60°C for three days (U40, Memmert GmbH & Co. KG, Germany). Subsequently, their initial dry mass, including body lipids, was measured (Sartorius M-Pact AX224), after which lipids were extracted using a petroleum ether extraction method [30, 38]. For this, dried specimens were placed in 5 mL petroleum ether for five days. Afterwards, the ether containing lipids was discarded and specimens were rinsed with fresh ether. The specimens were re-dried for another three days and weighed again post-extraction, allowing us to calculate the difference between the initial and post-extraction dry mass. The relative lipid content, serving as a proxy for fat body size and energy reserves, was calculated as absolute lipid content divided by initial dry mass.

### 2.6 Statistics and reproducibility

All statistical analyses and data visualizations were performed in R version 4.5.1 [39]. For full transparency and reproducibility, the complete code as including model selection procedures (provided as R Markdown output) and the dataset are available in the Zenodo Repository (https://doi.org/10.5281/zenodo.18298098). Experimental reproducibility was ensured by balancing initial sample sizes (pupae at the start of the experiment) across treatments and colonies (Table 1). Specifically, we used individuals from 5-6 colonies per treatment group, while naturally developed individuals were taken from 10 sources colonies (including four additional colonies), defining colonies as the primary unit of biological replication. We accounted for colony variability in all statistical models described in detail below.

To analyse the effect of cyclic thermal stress, while accounting for the effects of in vitro rearing by including data from colony-reared individuals where possible, and additional explanatory variables (e.g. sex, initial pupal weight) and their interactions on adult emergence rates, adult morphological traits (antennae and wing deformation, emergence weight, and head width), and relative lipid content, we fitted generalized linear mixed effect models (GLMMs) using the *glmmTMB* package [40]. Emergence rates and deformation status were fitted with binomial distribution (logit link), while the continuous variables proportional pupal weight loss, emergence weight, head width, and relative lipid content were fitted with a gamma distribution (log link). To analyse the duration of pupal development (number of days from the start of pupation to adult emergence), we applied an ordinal cumulative link mixed model (CLMM) with the Laplace approximation using the *ordinal* package [41], as assumptions for fitting a GLMM were not met (described below). While the CLMM included only colony ID as a random factor, GLMMs utilising only data from the *in vitro* rearing also included collection date as a random factor to account for colony and temporal-specific variability.

For each response variable, a set of candidate models was compared. The most parsimonious and best-fitting model was selected based on the Akaike information criterion (AIC), with lower AIC values indicating a better fit. The best-fitting models were compared to a simpler nested model using likelihood ratio tests (LRTs). When added terms did not improve model fit significantly, the simpler model was chosen and compared against its respective null-model using LRT. Model assumptions and dispersion of the data were checked using the *DHARMa* package [42]. When model assumptions were met, statistical significance (p < 0.05) was determined using the *Anova* function. Pairwise comparisons between treatment groups were conducted using the *emmeans* package [43] with Holm-Bonferroni corrections. Smooth predictions of emergence rate across the observed range of larval weights were generated using a prediction dataset with 100 evenly spaced values.

Differences in survival probability between treatment groups were initially determined for workers and males separately using Kaplan-Meier analyses via the *survival* and *ggsurvfit* packages [44], following to previous research [45]. To further analyse the main factors affecting adult survival, we fitted a set of candidate Cox Proportional Hazards (CoxPH) models using the *coxme* package [44]. The full model contained treatment (including data from individuals that developed naturally), sex (and their interactions), adult dry mass, relative lipid content, and deformations (antennae and wing deformation combined). The colony ID was included as a random factor. Model selection followed the AIC and LRT procedure as described above. The final best-fitting model to predict mortality included the explanatory variables treatment, sex, and their interactions, alongside deformation status. As proportional hazards (PH) assumptions were not entirely met (based on diagnostics using the *ggcoxdiagnostics* function of the *surmiver* package [46]), the final model was stratified by sex. Pairwise comparisons between treatment groups were performed using the *glht* function with Holm-Bonferroni correction.

## 3. Results

### 3.1 Emergence success and developmental duration

Cyclic thermal stress and pupal weight significantly affected the proportion of pupae reaching adulthood (pupal survival; GLMM: *χ*^2^ = 35.187, df = 3, p < 0.0001; Fig. 1*a,b*). This effect was primarily driven by the increase in thermal stress (*χ*^2^ = 22.276, df = 2, p < 0.0001; Fig. 1*a*). Pairwise comparisons revealed that pupae exposed to 37°C and 38°C cyclic thermal regimes had a 15% and 30% lower adult emergence rate, respectively, compared to pupae reared at constant 34°C (37°C vs 34°C: p < 0.05; 38°C vs 34°C: p < 0.0001). The difference between the 38°C and 37°C cyclic treatments was also significant (37°C vs 38°C: p < 0.05).

Emergence probability was significantly higher with an increase in initial pupal weight (*χ*^2^ = 11.065, df = 1, p < 0.001; figure 1*b*). Despite the marginally significant effect of sex on proportional pupal weight loss (GLMM: *χ*^2^ = 3.777, df = 1, p = 0.052; Fig. 1*d*, absolute pupal weight loss data are provided in figure S3*b*), including treatment did not improve model fit (LRT: *χ*^2^ = 0.904, df = 2, p = 0.904). Additionally, there was a strong positive relationship between adult emergence weight and the initial pupal weight (GLMM: *χ*^2^ = 402.03, df = 1, p < 0.0001; Pearson’s correlation: r = 0.88, p < 0.0001; Fig. S3*c*) with a small but significant effect of sex (*χ*^2^ = 4.03, df = 1, p = 0.045), whereas the interaction between initial pupal weight and sex had no effect (*χ*^2^ = 2.169, df = 1, p = 0.141). Including treatment did not significantly improve model fit (LRT: *χ*^2^ = 6.205, df = 3, p = 0.102)

While we observed a significant overall effect on pupation duration (CLMM: *χ*^2^ = 43.958, df = 3, p < 0.0001; Fig. 1*c*), this effect was not related to the thermal stress treatment (*χ*^2^ = 1.6121, df = 2, p = 0.447), but was sex-dependent (*χ*^2^ = 34.311, df = 1, p < 0.0001). Pairwise comparisons revealed that workers across all treatment groups were significantly different from males of all treatment groups (p < 0.001).

### 3.2 Carry-over effects on adult longevity

Cyclic thermal stress exposure during pupation resulted in reduced adult survival (longevity) for both workers (Log-rank test: *χ*^2^ = 8.9, df = 2, p = 0.01; Kaplan-Meier survival curves in Fig. 2*a*) and males *(χ*^2^ = 11.1, df = 2, p < 0.01; Fig. 2*b*). The presence of carry-over effects on longevity was strongly supported by the CoxPH model stratified by sex (*χ*^2^ = 517.34, df = 4, p < 0.0001, model concordance = 0.64 ± 0.01 SE; Fig. 2*e*). Pupae exposed to 38°C cyclic treatment had a significantly higher mortality risk as adults than pupae reared at 34°C (HR = 2.00, 95% CI = 1.61-2.48, p < 0.0001, Fig. 2*e*). The 37°C cyclic treatment during pupal development posed a marginally significant higher mortality risk during adulthood (HR = 1.39, 95% CI = 1.024-1.89, p = 0.035, Fig. 2*e*). We also found that individuals with morphological deformations (combining deformed antennae and wings) had a significantly higher mortality risk (HR = 2.38, 95% CI = 1.94-2.92, p < 0.0001, Fig. 2*e*). Importantly, we found no evidence that mortality risk during adulthood significantly differed between individuals reared *in vitro* as L4 larvae and pupae at 34°C and those developed naturally in their natal colony until adult emergence (HR = 0.95, 95% CI = 0.70-1.30; p = 0.756, Fig. 2*e*). This was consistent with the survival analyses in both adult workers (Log-rank test: *χ*^2^ = 0, df = 1, p = 0.9, Fig. 2*c*) and males (*χ*^2^ = 0.8, df = 1, p = 0.4, Fig. 2*d*).

**Figure 2.**
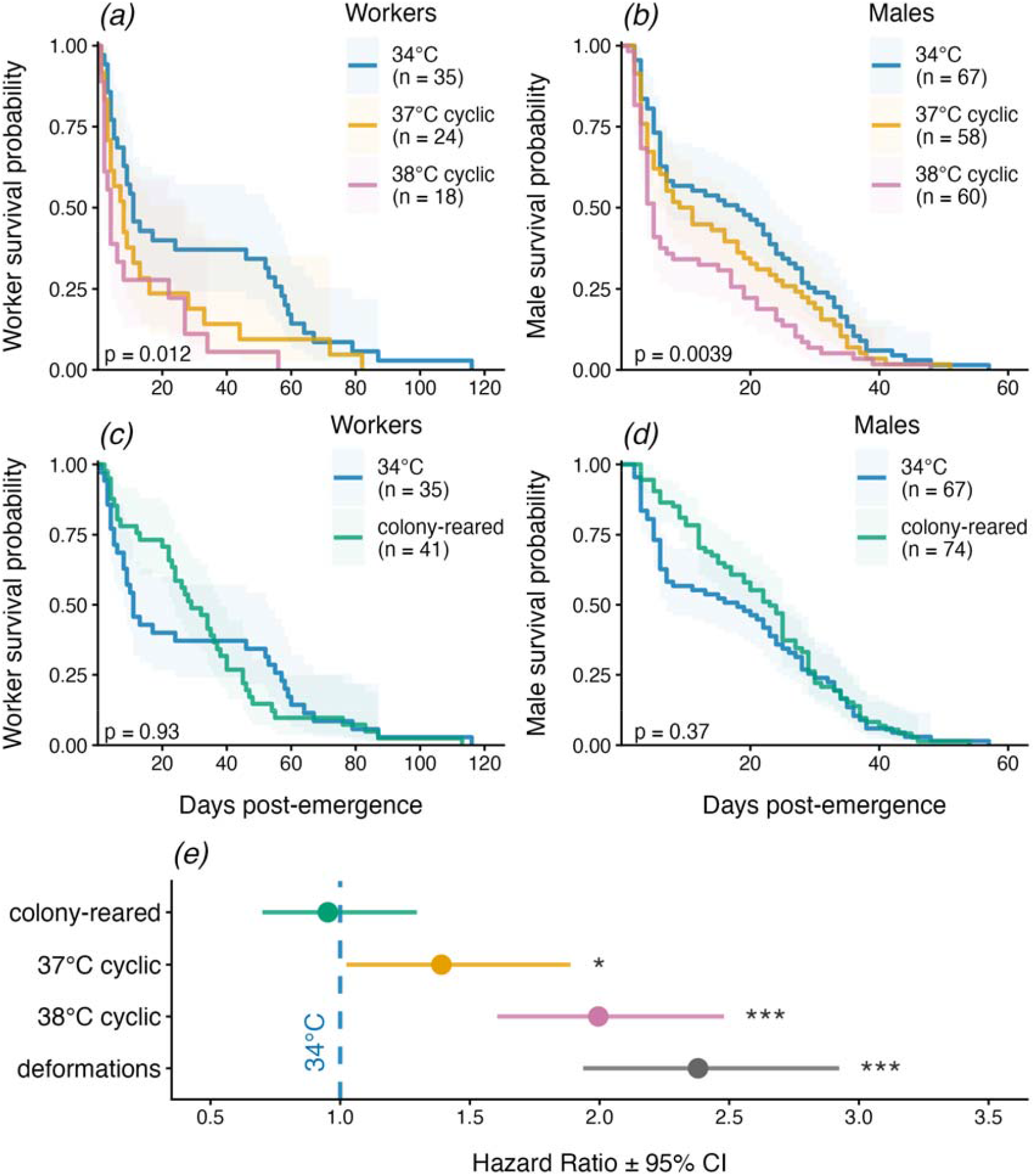
Carry-over effects of cyclic thermal stress during pupation on adult longevity in *B. terrestris*. Kaplan-Meier survival curves (± 95% CI, shaded areas) showing the carry-over (indirect) effect of thermal treatments during pupation *in vitro* on the longevity in adult workers **(*a*)** and males **(*b*)**. The group kept at a constant 34°C (blue) is compared to the cyclic thermal stress treatments (37°C cyclic, orange; 38°C cyclic, reddish-purple). Comparison of *in vitro* rearing of L4 larvae and pupae at 34°C (blue) to those naturally developed in their natal colony (bluish green) until emergence on adult survival in workers **(*c*)** and males **(*d*)**, respectively. Sample sizes (n) and overall statistical significance (p) from the Kaplan-Meier analyses are provided in each plot. **(*e*)** Forest plot of the hazard ratios (HR) ± 95% CI based on the stratified CoxPH model for treatment (cyclic thermal stress during pupal development) and morphological deformations (combined antennae and wings deformations; Fig. S2) on adult longevity. HR values > 1 indicate significantly increased hazard. Asterisks denote significance (* p < 0.05; *** p < 0.001) relative to the 34°C group (dashed blue line).

### 3.3 Morphology and energy reserves

We found evidence that cyclic thermal stress during pupal development increased the incidence of deformed wings in adults (GLMM: *χ*^2^ = 20.383, df = 3, p = 0.0001; Fig. 3*a*, filled bars; Fig. S2*b*). Pupae exposed to cyclic 38°C developed deformed wings more frequently than pupae kept at 34°C (38°C vs 34°C: 3.6-fold increase, p < 0.01) and those reared inside their natal colony (38°C vs colony: 4.5-fold increase, p < 0.001), while there was no significant difference compared to pupae exposed to cyclic 37°C (38°C vs 37°C: p = 0.155). Nonetheless, there was no significant difference between pupae exposed to cyclic 37°C compared to pupae kept at 34°C (37°C vs 34°C: p = 0.294) or those reared inside their natal colony (37°C vs colony: p = 0.169). There was no significant difference in the incidence of deformed wings between pupae reared *in vitro* at 34°C compared to colony-reared individuals (34°C vs colony: p = 0.636). Furthermore, there was no significant effect of thermal treatment during pupal development on the incidence of adult bees with deformed antennae (*χ*^2^ = 6.241, df = 3, p = 0.100; Fig. 3*a*, open bars; Fig. S2*a*).

**Figure 3.**
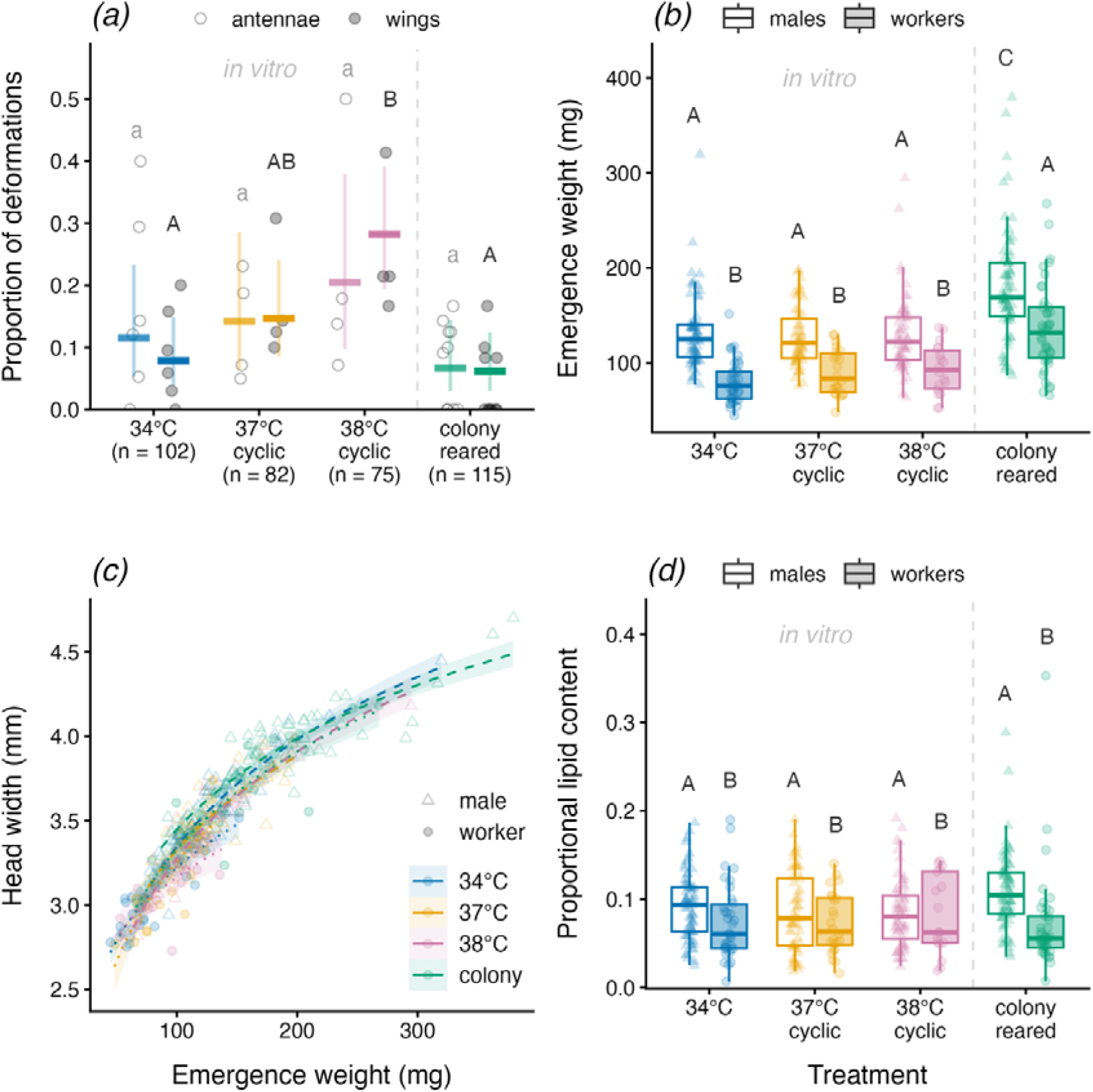
Effects of cyclic thermal stress during pupation on morphological and physiological traits in adult *B. terrestris*. ***(a)*** Proportion of individuals per colony (n ≥ 5 per colony) with deformed antennae (open circles; Fig. S2*a*) and wings (filled grey circles; figure S2*b*), and the GLMM means ± 95% CI (bars). Sample sizes (n) are provided below for each treatment group. Colours **(*a-d*)** indicate different rearing conditions *in vitro* (treatments during pupal development: blue = 34°C; orange = 37°C cyclic; reddish-purple = 38°C cyclic) and inside their natal colony (bluish green). **(*b*)** Box plots showing the weight of newly emergence adults (males = open boxes; workers = filled boxes) for each treatment. **(*c*)** Scaling of adult head width relative to emergence weights for adults from each treatment group. Lines (dashed = males; dotted = workers) show fitted trends ± 95% CI from GLMs (ignoring colony effects) with log-transformed emergence weight. **(*d*)** Box plots showing the relative lipid content (proxy for remaining energy reserves) in adults at the end of their lifespan for each treatment group. Individual data points in **(*b-d*)** are shown as triangles (males, m) and circles (workers, w). Sex-specific samples sizes for 34°C (n_m_ = 67, n_w_ = 35), 37°C cyclic (n_m_ = 58, n_w_ = 24), 38°C cyclic (n_m_ = 60, n_w_ = 18), and colony-reared (n_m_ = 74, n_w_ = 41). Significant differences (p < 0.05), based on post-hoc comparisons from GLMMs, are indicated by different letters above the bars and boxes (groups sharing a letter are not significantly different).

We found a significant effect on adult emergence weight (GLMM: *χ*^2^ = 158.96, df = 4, p < 0.0001; Fig.3*b*), which was both treatment-dependent (*χ*^2^ = 82.129, df = 3, p < 0.0001) and sex-dependent (*χ*^2^ = 94.806, df = 1, p < 0.0001) without an interaction effect. Pairwise comparisons revealed that the emergence weight of adults reared *in vitro* during larval and pupal development was lighter compared to adults that emerged in their natal colony (colony vs all *in vitro* treatments: p < 0.0001), and males were significantly heavier than workers across all treatment groups (males vs workers: p < 0.0001).

Further morphometric analyses revealed that head width strongly increased with emergence weight (GLMM: *χ*^2^ = 1403.577, df = 1, p < 0.0001; Fig. 3*c*). This relationship was also both treatment-dependent (*χ*^2^ = 50.95, df = 3, p < 0.0001) and sex-dependent (*χ*^2^ = 40.242, df = 1, p < 0.0001) with a weak three-way interaction (weight*treatment*sex: *χ*^2^ = 15.966, df = 3, p < 0.001), but no significant two-way interactions (weight*treatment: *χ*^2^ = 2.102, df = 3, p = 0.552, weight*sex: *χ*^2^ = 0.417, df = 1, p = 0.518, treatment*sex: *χ*^2^ = 0.861, df = 3, p = 0.835). However, the scaling relationship between emergence weight and head width remained largely consistent across treatments and sexes, with slopes ranging from 0.17 to 0.27 and no significant differences between slopes (all interactions p > 0.05, detailed comparisons in Table S1; Fig. 3*c*).

There was also a strong morphometric relationship between head width and intertegular distance (ITD) measured in adults (GLMM: *χ*^2^ = 567.94, p < 0.0001; Pearson’s correlation: r = 0.87, Fig. S3*d*), with similar scaling relationships between sexes (both slopes = 0.353) despite males being larger than workers (*χ*^2^ = 5.43, p < 0.05). Including treatment did not significantly improve the model fit (LRT: *χ*^2^ = 6.205, p = 0.102).

Analysis of relative lipid content (as a proxy for energy reserves) at the end of adult life revealed a significant effect of sex (GLMM: *χ*^2^ = 4.27, df = 1, p = 0.039, Fig. 3*d*), which was independent of treatment, as including treatment did not improve the model fit (LRT: *χ*^2^ = 0.923, df = 2, p = 0.631).

## 4. Discussion

Our study addresses the knowledge gap concerning the lethal and sublethal, long-term carry-over effects of thermal extremes in insect pollinators [3, 4, 10]. We provide empirical evidence that thermal stress during pupal development leads to deferred mortality at the adult life-stage in bumblebees. This finding reveals a mechanism of deferred mortality that may contribute significantly to pollinator decline. Specifically, cyclic thermal stress not only caused acute pupal mortality (i.e. reduced emergence; Fig. 1*a*) but also resulted in significantly reduced adult longevity (HR = 2.00) in both workers and males of *B. terrestris* (Fig. 2*a,b,e*). This finding suggests that thermal stress during pupation imposes irreversible developmental damage that undermines adult physiological resilience.

### Developmental bottlenecks and irreversible damage

The high sensitivity observed during the pupal phase underscores its role as a critical developmental bottleneck. Reductions in emergence success by 15% and 30% under cyclic thermal stress of 37°C and 38°C respectively (Fig. 1*a*) align with patterns observed under constant thermal stress [30], though to a lesser extent. Noteworthy, average emergence rates of about 80% at 34°C were slightly higher than in our previous *in vitro* study under constant thermal stress [30] and for *B. impatiens* colonies exposed to 25°C, 30°C, and 35°C [23], likely reflecting temporal, colony, and species variation. Interestingly, despite high emergence rates reported for those *B. impatiens* colonies, there was also a drastic reduction in the number of offspring for colonies housed at 35°C, indicating colony failure which could be linked to the increase in adult workers abandoning their nest [23]. This observation may indicate insufficient nursing (feeding) of larvae at 35°C. Indeed, our observation that heavier pupae (strongly dependent on L4 larvae weight, Fig. S3*a*) were more likely to emerge as adults (Fig. 1*b*) supports the idea that successful pupa-to-adult transition might be resource-dependent [30]. This suggests that resource limitations within a colony could trigger negative synergistic interactions, amplifying the detrimental effects of thermal stress.

We found no significant effect on the duration of metamorphosis (pupation) between the cyclic thermal stress treatments and incubating pupae at a constant 34°C (Fig. 1*c*). Regardless, pupation duration differed between workers (8-9 days) and males (9-10 days), aligning with data reported for pupae reared in *B. impatiens* colonies [47] and *B. terrestris* reared *in vitro* [30, 37]. In contrast to how constant thermal stress of 38°C may slightly prolong pupal development, exposure to 36°C appears to shorten developmental time compared to 34°C [30].

Most notably, 38°C cyclic thermal stress led to a higher incidence of deformed wings (Fig. 3*a*), but this effect was much less pronounced compared to pupae exposed to a constant temperature of 38°C for five days during pupation [30]. Interestingly, cold stress during pupal development, mimicking a spring cold snap, has been shown to cause wing deformities and reduced flight ability in the alfalfa leafcutting bee *Megachile rotundata* (Hymenoptera: Apiformes: Megachilidae) [33]. While wing morphology is a known indicator of irreparable developmental damage [30, 48, 49], our data provide clear empirical evidence that deformations (antennae and wings) can serve as a powerful hazard indicator (HR = 2.38) that predicts adult longevity regardless of treatment (Fig. 2*e*). Although survival was reduced in adults that were exposed to the cyclic 38°C treatment (Fig. 2*a,b,e*), there was a steep decline in adult survival during the first two weeks. We speculate that this may stem from a lack of social interactions [50]. Supporting this and validating our *in vitro* rearing approach, no significant difference was detected in the mortality risk during adulthood between individuals developed naturally in their natal colonies until emergence and those reared *in vitro* at 34°C as L4 larvae and pupae (HR = 0.95; Fig. 2*c,d*).

Beyond individual mortality, workers and males with severely deformed wings are unable to fly, which consequently reduces colony fitness. In fact, it has already been shown that thermal stress during development can reduce adult fitness across diverse insect taxa [6, 7, 9]. Adding to the consequences of deformed wings, deformed antennae likely impair the ability to detect scents of nestmates, mating partners (important for males), and floral resources (important for workers). This idea is supported by a study demonstrating heat stress-induced olfactory impairment in adult bumblebee workers [51].

### Decoupling fitness costs from adult morphometrics and fat body

Despite the difference in functional longevity, there was no evidence that deferred mortality was associated with easily measurable morphometric traits (i.e. pupal weight loss, emergence weight, head width, and ITD) between *in vitro* treatments (Fig. 3*b-d* and S3*b-d*). While this result suggests that selection pressure from short-term thermal stress for body size might be more pronounced at earlier larval stages [30, 31], this is in contrast with some previous colony-level findings. For example, a smaller male wing size was reported for *B. terrestris* micro-colonies housed at 33°C compared to 26°C [26]. Furthermore, worker offspring were shown to have a smaller ITD in colonies housed at 33°C compared to 25°C, whereas males appeared to be unaffected [25, 27]. It is worth noting that our colony-reared individuals, housed at 25°C, also emerged slightly heavier than *in vitro* reared bees (Fig. 3*b*). Although this difference could indicate a temperature-related ceiling effect, with pupae reared at 34°C *in vitro* being already heat-stressed, our chosen baseline aligns with brood temperatures measured in *B. terrestris* nests [19, 28, 29].

An alternative explanation for our observed weight difference would be the possibility of initial food consumption by newly emerged bees inside the nest, unlike bees emerging from the *in vitro* rearing. However, as the median proportional pupal weight loss was almost identical between treatments (Fig. 1*d* and S3*b*) with a strong positive relationship between initial pupal weight and emergence weight (Fig. S3*c*) and the scaling between emergence weight and head width was also very similar across all treatment groups (Fig. 3*c*), we think that our observed pattern reflects the limitations of *in vitro* rearing in general compared to naturally reared larvae inside the colony. It is very likely that colony-reared L4 larvae received more food than our *in vitro* reared L4 larvae, as workers can sense hungry larvae and provide food whenever needed [52].

We would even speculate that the body size effects observed in thermally stressed colonies of the aforementioned studies [25-27] play out at an earlier larval stage [30, 31] and are an indirect result of altered worker nursing and foraging behaviour [23, 24] rather than a direct physiological consequence of the heat itself. It could well be that either resources are much more limited or larvae are simply insufficiently fed by thermally stressed adult workers. This highlights the importance of using an *in vitro* approach to isolate direct physiological effects from comprehensive colony dynamics [30, 31].

While increased lipid content is advantageous for constantly heat-stressed pupae to successfully emerge [30], we found no significant difference in relative lipid content at the end of their adult life (Fig. 3d), aligning with previous findings of energy homeostasis in ageing bumblebee workers [53]. This indicates the capacity of surviving adults to maintain homeostasis of their fat body, which is the primary site for heat shock protein (HSP) synthesis and storage of energy reserves [48, 54, 55], by resource allocation or compensatory foraging. Hence, any sublethal damage carrying over to adulthood might be genetically, structurally, and/or physiologically mediated rather than simply due to depleted energy reserves. This aligns with findings where developmental thermal stress indirectly affected insect adult physiology, such as flight performance and behaviour [56-58].

## Conclusion

These findings highlight that sublethal thermal stress during insect development may lead to profound fitness costs. Our study provides compelling empirical evidence that cyclic thermal stress during pupal development significantly reduces adult longevity in *B. terrestris*. Furthermore, our findings confirm that 38°C is a critical threshold for developmental disruption in *B. terrestris* [30, 31]. Although our study oversimplifies realistic thermal stress scenarios, such as those experienced during heatwaves, and *B. terrestris* only serves as a model system, we are convinced that these findings provide valuable insights into the consequences of thermal stress during pupal development in bumblebees. Species that are less heat-tolerant than *B. terrestris*, such as *B. lapidaries* and *B. alpinus* [13], or typically nest above ground, such as *B. hypnorum* and *B. pascuorum* [19] or are used in greenhouses for pollination services, would be expected to be especially at risk [30]. As sublethal carry-over effects are rarely considered, current prediction models may underestimate the long-term impact of environmental stressors on crucial pollinator populations. To generate a more holistic picture of environmental stress impact, gaining a nuanced mechanistic understanding of carry-over effects across diverse insect species and life stages, incorporating both individual-level, and for social insects, also colony-level physiological and life-history data, remains crucial.

## Data accessibility statement

Dataset and R script used for statistical analysis and producing all Fig. are available from the Zenodo Repository: https://doi.org/10.5281/zenodo.18298098. Electronic supplementary material is available online.

## Statement of authorship

M.M.M.: methodology, investigation, data curation. C.K.: conceptualization, methodology, investigation, data curation, resources, validation, formal analysis, writing – original draft preparation, writing – review and editing, visualization, supervision, project administration.

## Acknowledgments

We thank Sandra Laußer, Laura Wögler, Ramona Schöpf, and Michael Ott for their assistance in the laboratory. We are grateful to Prof. Dr. Tomer J. Czaczkes for sponsoring us bumblebee colonies. We have used AI for proofreading.

## Ethics statement

This study was conducted in accordance with the ethical regulations of the German Animal Welfare Act (TierSchG) for conducting experiments with insects.

