## Supplementary Figure S1-3 and Table S1 for "Deferred mortality: cyclic thermal stress during pupation triggers irreversible carry-over effects in a key pollinator"

### Electronic supplementary material

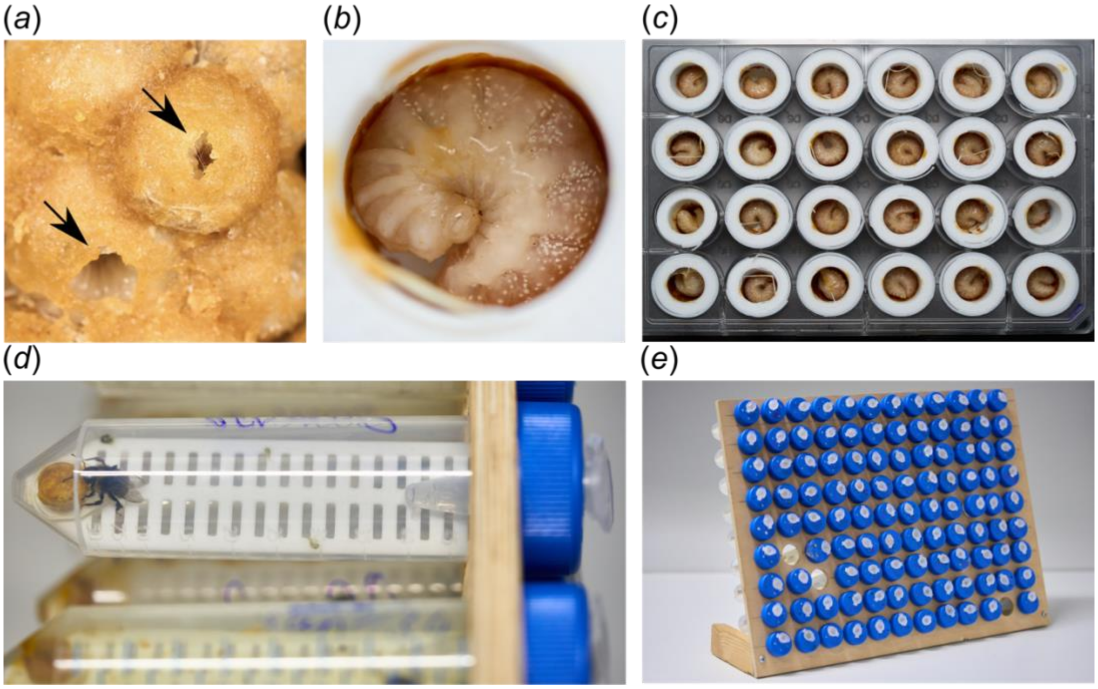


**Figure S1. Experimental setup for *in vitro* rearing and adult longevity assay in *B. terrestris*.** (*a*) Fourth-instar larvae (L4) were identified by the characteristic small feeding holes (black arrows), collected, and transferred into (*b*) 3D-printed polylactide (PLA) artificial brood cells for *in vitro* rearing (see methods). These cells were within (*c*) a 24-well plate setup inside humidity and temperature controlled incubators. (*d*) Newly emerged adults were individually housed in modified 50 mL tubes, serving as individual cages. Each caged contained a 3D-printed PLA mesh insert for sanitary reasons, a 1.5 mL perforated tube providing *ad libitum* 60% w/v sucrose solution, and pollen candy. (e) Individual cages were randomly stored in a wooden rack within the insectary room of their natal colonies, maintained at standardized conditions (25 ± 1°C and 30-50% RH).

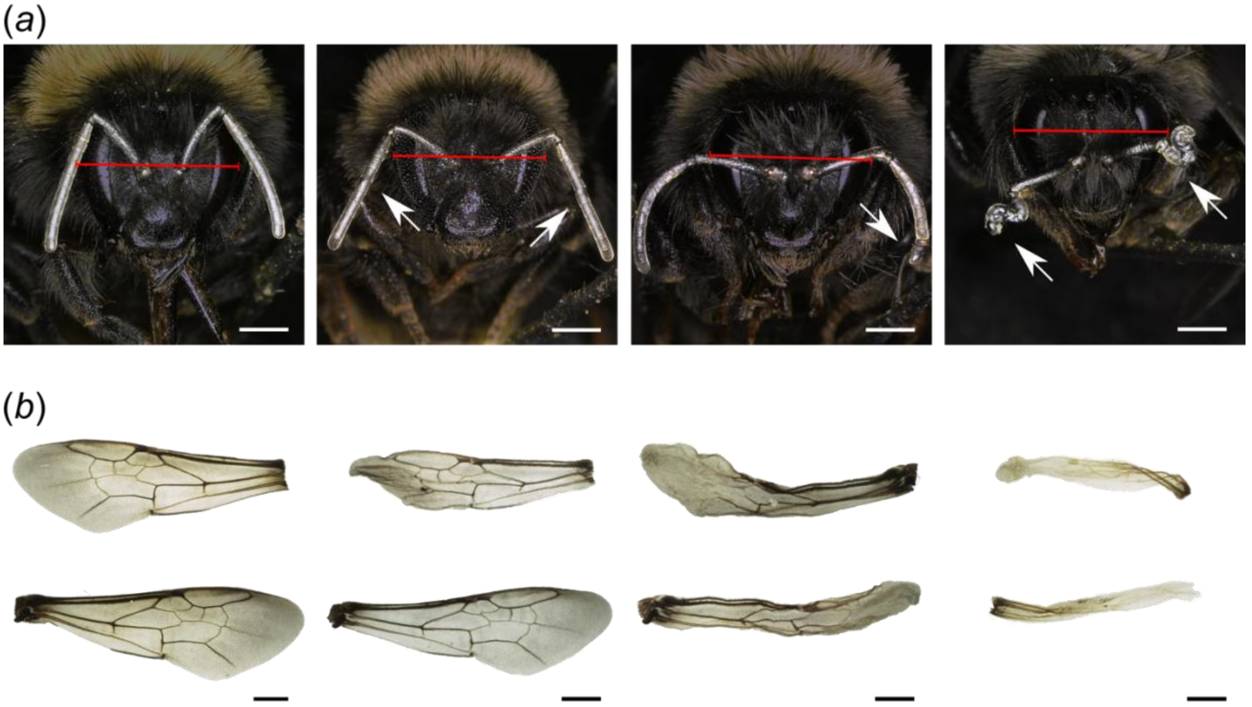


**Figure S2. Representative examples of morphological deformations in adult *B. terrestris*.** (*a*) Examples ranging from an individual with normally developed antennae (left) to increasingly severe deformations (right). White arrows indicate regions of deformations, including slightly deformed, fused, missing and abnormally curved flagellomeres. Red lines indicate head width measurements. (*b*) Paired left and right forewings from an adult bee with normal development (left) to examples with increasingly more severe deformations (right). These images serve as representative examples of what we scored as deformations in our analyses (see methods; Fig. 2*e* and 3*a*). For the presentation purposes, photographs were enhanced by colour corrections (heads and wings), shading (heads), removing backgrounds, and aligning (wings) using graphical software. Scale bars = 1mm.

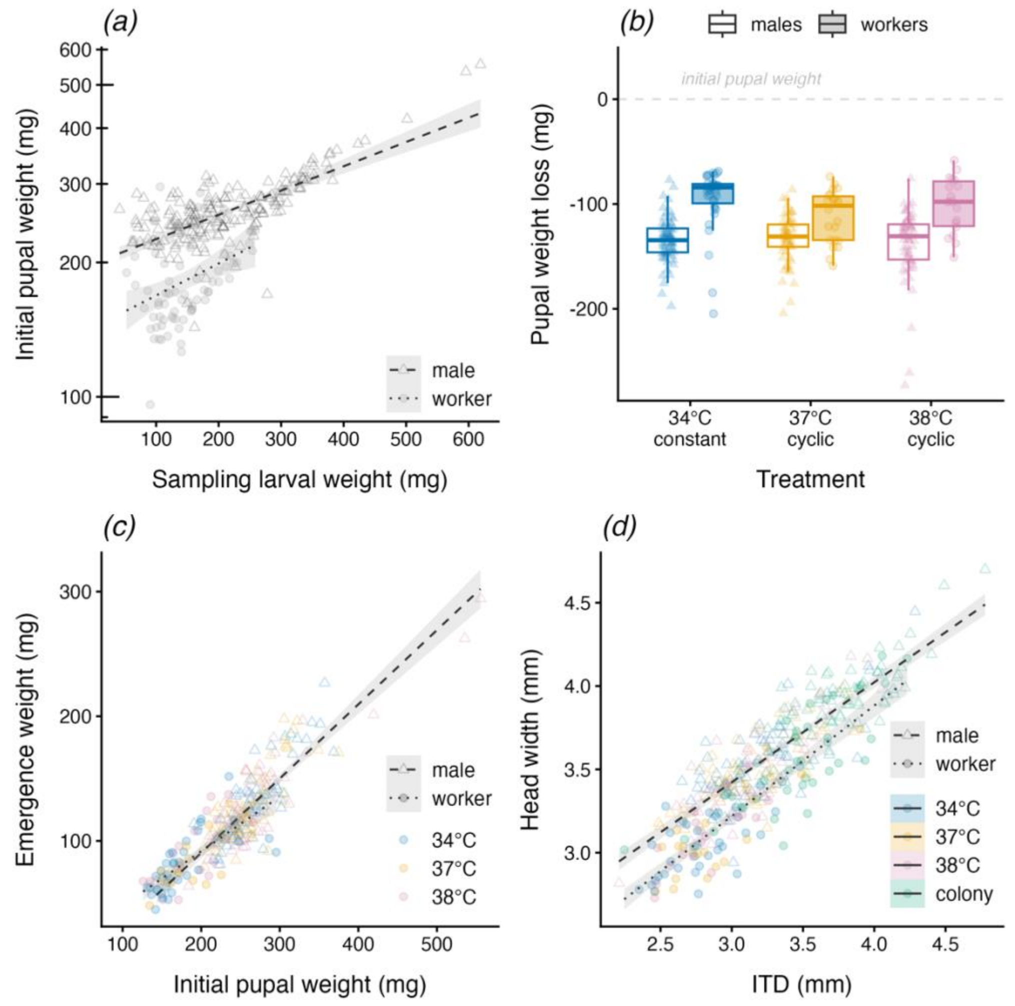


**Figure S3. Morphometric relationships from initial sampling of L4 larvae to adulthood in *B. terrestris* males and workers.** ***(a)*** There was a strongly positive relationship between the sampling weight (mg) of L4 larvae from their natal colonies and their initial pupal weight (on log10 scale) at the beginning of the experiment (lognormal GLMM: $\chi^2$ = 260.65, $p < 0.0001$), with pupal weight differing between sexes ($\chi^2$ = 110.82, $p < 0.001$). ***(b)*** Box plots show absolute weight loss during pupation (initial pupal weight – weight at emergence) for each *in vitro* rearing treatment and sex (open boxes = males; filled boxes = workers). The proportional weight loss (relative to initial weights) and statistical analysis are presented in the main manuscript (see Fig. 1*d*). ***(c)*** There was strong positive linear relationship between initial pupal weight (mg) and emergence weight (mg) (GLMM: $\chi^2$ = 402.03, $p < 0.0001$; Pearson's correlation: r = 0.88), which was independent of treatment (blue = constant 34°C; orange = cyclic 37°C; reddish-purple = cyclic 38°C), but a marginally sex-dependent effect ($\chi^2$ = 4.03, $p = 0.045$). ***(d)*** Morphometric relationship between ITD (inter tegular distance) and head width measured in adults of colony-reared (green = colony) and *in vitro*-reared individuals (i.e. experimental treatments: blue = constant 34°C; orange = cyclic 37°C; reddish-purple = cyclic 38°C). ITD was a strong predictor of head width (GLMM: $\chi^2$ = 567.94, $p < 0.0001$; Pearson's correlation: r = 0.87), with indistinguishable scaling relationships between sexes (both slopes = 0.353), but with males being larger than workers ($\chi^2$ = 5.43, $p < 0.05$). Individual data points in all plots are displayed as triangles (males) and circles (workers).

**Table S1. Pairwise comparisons of slopes describing the scaling relationship between emergence weight and head width across treatments and sexes.** Estimates are derived from the GLMM including a significant three-way interaction between emergence weight, treatment (34 °C constant, 37 °C cyclic, 38 °C cyclic, and colony-reared), and sex (males and workers). Despite this interaction, scaling relationships remained largely consistent across all groups (see Fig. 3c).

| Pairwise comparisons | Estimate | SE | df | z ratio | p value |
|---|---|---|---|---|---|
| 34°C male – 37°C male | 0.0193 | 0.0239 | Inf | 0.8105 | 1.0000 |
| 34°C male – 38°C male | -0.0088 | 0.0215 | Inf | -0.4122 | 1.0000 |
| 34°C male – colony male | 0.0442 | 0.0202 | Inf | 2.1819 | 0.6406 |
| 34°C male – 34°C worker | 0.0440 | 0.0256 | Inf | 1.7202 | 1.0000 |
| 34°C male – 37°C worker | -0.0210 | 0.0300 | Inf | -0.6985 | 1.0000 |
| 34°C male – 38°C worker | 0.0829 | 0.0304 | Inf | 2.7266 | 0.1664 |
| 34°C male – colony worker | 0.0122 | 0.0219 | Inf | 0.5571 | 1.0000 |
| 37°C male – 38°C male | -0.0282 | 0.0235 | Inf | -1.1969 | 1.0000 |
| 37°C male – colony male | 0.0248 | 0.0225 | Inf | 1.1059 | 1.0000 |
| 37°C male – 34°C worker | 0.0246 | 0.0272 | Inf | 0.9054 | 1.0000 |
| 37°C male – 37°C worker | -0.0403 | 0.0313 | Inf | -1.2868 | 1.0000 |
| 37°C male – 38°C worker | 0.0636 | 0.0319 | Inf | 1.9898 | 0.9150 |
| 37°C male – colony worker | -0.0071 | 0.0239 | Inf | -0.2986 | 1.0000 |
| 38°C male – colony male | 0.0530 | 0.0200 | Inf | 2.6534 | 0.1992 |
| 38°C male – 34°C worker | 0.0528 | 0.0252 | Inf | 2.0955 | 0.7586 |
| 38°C male – 37°C worker | -0.0121 | 0.0297 | Inf | -0.4085 | 1.0000 |
| 38°C male – 38°C worker | 0.0918 | 0.0302 | Inf | 3.0405 | 0.0661 |
| 38°C male – colony worker | 0.0210 | 0.0216 | Inf | 0.9762 | 1.0000 |
| colony male – 34°C worker | -0.0002 | 0.0242 | Inf | -0.0081 | 1.0000 |
| colony male – 37°C worker | -0.0651 | 0.0290 | Inf | -2.2470 | 0.5667 |
| colony male – 38°C worker | 0.0387 | 0.0294 | Inf | 1.3185 | 1.0000 |
| colony male – colony worker | -0.0320 | 0.0203 | Inf | -1.5755 | 1.0000 |
| 34°C worker – 37°C worker | -0.0650 | 0.0325 | Inf | -1.9977 | 0.9150 |
| 34°C worker – 38°C worker | 0.0389 | 0.0332 | Inf | 1.1718 | 1.0000 |
| 34°C worker – colony worker | -0.0318 | 0.0256 | Inf | -1.2420 | 1.0000 |
| 37°C worker – 38°C worker | 0.1039 | 0.0368 | Inf | 2.8257 | 0.1274 |
| 37°C worker – colony worker | 0.0332 | 0.0301 | Inf | 1.1023 | 1.0000 |
| 38°C worker – colony worker | -0.0707 | 0.0305 | Inf | -2.3213 | 0.4864 |
